# Agent-based simulation of large tumors in 3-D microenvironments

**DOI:** 10.1101/035733

**Authors:** Ahmadreza Ghaffarizadeh, Samuel H. Friedman, Paul Macklin

## Abstract

Multicellular simulations of tumor growth in complex 3-D tissues, where data come from high content *in vitro* and bioengineered experiments, have gained significant attention by the cancer modeling community in recent years. Agent-based models are often selected for these problems because they can directly model and track cells’ states and their interactions with the microenvironment. We describe PhysiCell, a specific agent-based model that includes cell motion, cell cycling, and cell volume changes. The model has been performance tested on systems of 10^5^ cells on desktop computers, and is expected to scale to 10^6^ or more cells on single super-computer compute nodes. We plan an open source release of the software in early 2016 at PhysiCell.MathCancer.org

## I. INTRODUCTION

A detailed analysis of the interactions between tumor cells and their microenvironment is required for a comprehensive understanding of tumor development, growth, therapy response, and evolution of resistance to therapy. Structural and biochemical properties of the microenvironment, along with the dynamic features of the tumor cells that reside within and interact with it, can impact the complex process of metastasis. Thus, a realistic tumor model should take both cells and microenvironment into account simultaneously.

Over the past several years, a variety of models and tools have been developed to help simulate tumor growth and understand the various mechanisms involved in this process [1-3]. These methods are either general modeling frameworks that can be tailored to cancer studies (e.g., [4-7]), or models designed specifically for tumor simulations (e.g., [8-11]).

In this article, we discuss PhysiCell: a particular off-lattice agent based model, designed as a general modeling framework, but with cancer simulations in mind. We show how this tool models cell and microenvironment characteristics to simulate a 3-D tumor in which the cells interact with the other cells as well as the microenvironment. It is worth noting that the system does not fit or prescribe tumor growth or structure, but rather models basic conservation laws (e.g., diffusive substrate transport and consumption, and Newton’s second law of motion for tumor cells) and prescribes simple, biologically-driven rules of individual cell behaviors, from which emerge the observed tissue-scale tumor growth phenomena such as a viable rim and large central necrotic core. This is one of the key motivations for using mechanistic, agent-based models: to test the behaviors that emerge from basic biological and physical hypotheses, and evaluate model predictions against *in vitro* and other multicellular data.

With PhysiCell, it is possible to directly simulate 3-D *in vitro* experiments, allowing more direct model comparison, faster hypothesis testing, and expansion of *in vitro* insights to their probable *in vivo* and clinical impacts.

**Authors’ note:** This work describes early progress in the PhysiCell project. Please check PhysiCell.MathCancer.org for the most up-date project information and downloads.

## II. METHODS

### A. Approach

Tumor cells, like any other physical objects, obey the laws of physics. So, we start modeling a tumor by writing the equations that govern the physical behavior of the cells.

#### 1) Cell Motion

Cell motion is the result of balance between biomechanical forces exchanged with other cells and the microenvironment. The cell-cell forces differ from one cell type to another; for example, the cells undergoing epithelial-mesenchymal transition experience large changes in their adhesion markers [12]. Equation 1 shows the balance of forces acting on cell *i*. PhysiCell solves Equation 1 with an intertialess assumption (d/dt *m_i_v_i_* ≈ 0 as in [9, 13]) to calculate the cell *i* velocity.

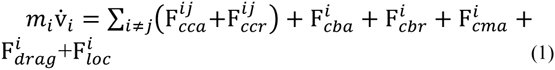

where F*_cca_* and F*_ccr_* represent cell-cell adhesion and repulsion, respectively, F*_cba_* and F*_cbr_* represent cell-basement membrane (if present) interactions (adhesion and repulsion respectively), F*_cma_* accounts for cell-ECM adhesion, and *F_drag_* is drag. Motile cells experience a net locomotive force (F*_loc_*).

#### 2) Cell cycling

PhysiCell gives each cell a phenotypic state S(*t*) at each time *t*, that can be C (cycling), A (apoptotic), or N (necrotic). Each state can have one or more phases. For example, C can have phases *G*_0_/*G*_1_, *S*, *G*_2_, and *M* with different time scales *τ*_G0/G1_, *τ*_S_, *τ*_G2_, *τ*_M_ respectively. Cells sample the environment and use the microenvironment parameters as the inputs when making decisions about changing their states. This sampling enables PhysiCell to incorporate experimental observations and theory into the models. For example, we know that if glucose or oxygen levels are low or the cell compression is high, then *τ*_G0/G1_ will be long (allowing us to model cell cycle arrest). So we can write this timescale as:

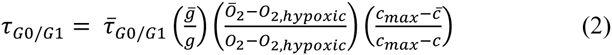

where *g* and *O_2_* are the concentration of glucose and oxygen in the microenvironment, and *c* is the cell's compression.

#### 3) Cell volume regulation

Cell volume changes during proliferation and death are key to analyzing tumor growth. PhysiCell uses a system of ODEs to model changes in the cytoplasmic and nuclear fluid volume (*V*_CF_, *V*_NF_) and cytoplasmic and nuclear solid biomass volume (*V*_CS_, *V*_NS_), with simple parameter changes to model phenotype changes during cycle progression or death:

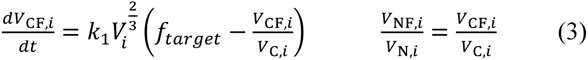

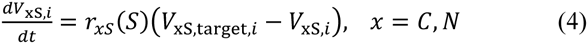

### B. Results

Figure 1 shows a simulated 3-D *in vitro* tumor spheroid, where we seeded 10 tumor cells in an 8 mm^3^ (2 mm × 2 mm × 2 mm) domain, where the oxygen distribution is solved with a reaction-diffusion solver [14]. The tumor in Figure 1 has over 557,000 cells that form a 1.7 mm necrotic tumor spheroid after 42 days of growth. In this figure, cycling cells are green, nuclei are blue, quiescent cells are pale blue, apoptotic cells are red, and necrotic ones are brown.

**Figure 1:**
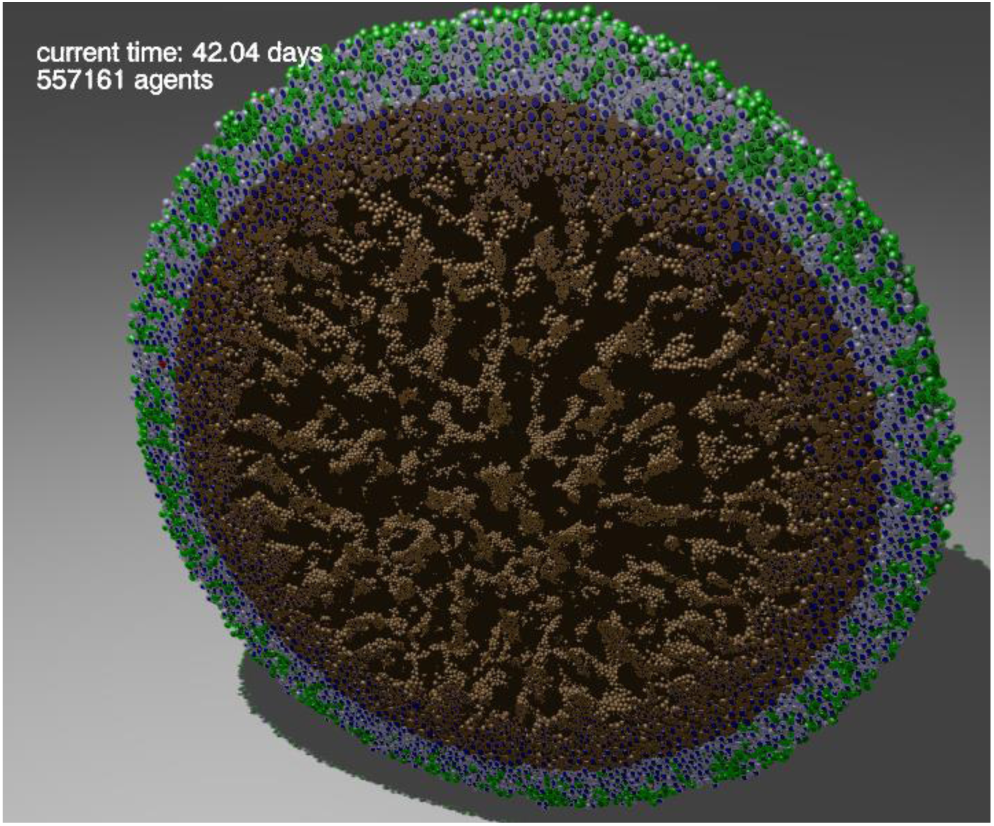
A 1.7 mm necrotic tumor spheroid containing over 557,000 cells. Cycling (green), quiescent (pale blue), and apoptotic cells form a viable rim. Nuclei are blue and necrotic cells are brown. *in vitro* tumor spheroids exhibit similar morphologies.

## III. CHALLENGES AND OPPORTUNITIES

Like any other agent based model, PhysiCell is limited by the number of cells (agents) and size of the microenvironment (with respect to the resolution of voxels) it can simulate. The efficient implementation of PhysiCell (including parallelization by OpenMP) enables it to scale the computing cost linearly with the cell count. PhysiCell is able to perform simulations with ~10^5^ cells on a regular desktop, and is expected to scale to 10^6^ cells on single supercomputer compute nodes.

PhysiCell uses the multicellular data standard (Multi-CellDS: see MultiCellDS.org) to read input and write outputs, with calibration to high-throughput screening data recorded as digital cell lines. MultiCellDS aims to create a standard data format for multicellular modeling frameworks. This will help for sharing experimental, simulation, and clinical data, and for comparing and ensembling independent models seeded with identical initial data.

## ACKNOWLEDGMENT

We thank the USC Center for Applied Molecular Medicine for generous resources. This work was supported by the Breast Cancer Research Foundation for support, the National Institutes of Health (Physical Sciences Oncology Center grant 5U54CA143907 for Multi-scale Complex Systems Transdisciplinary Analysis of Response to Therapy (MC-START), and 1R01CA180149), and the USC James H. Zumberge Research and Innovation Fund. We thank Margy Gunnar for her code testing contributions.

